# The impact and implications of vehicle collisions on the endangered and endemic Zanzibar red colobus (*Piliocolobus kirkii*)

**DOI:** 10.1101/2020.01.17.906909

**Authors:** Harry Olgun, Mzee Khamis Mohammed, Abbas Juma Mzee, M. E. Landry Green, Tim R. B. Davenport, Alexander V. Georgiev

**Affiliations:** School of Natural Sciences, Bangor University, & Zanzibar Red Colobus Project; Department of Forestry and Non-Renewable Natural Resources, Revolutionary Government of Zanzibar, Zanzibar, Tanzania; School of Natural Sciences, Bangor University, Bangor, UK & Zanzibar Red Colobus Project; Wildlife Conservation Society (WCS), Zanzibar, Tanzania

**Keywords:** colobus, Jozani-Chwaka Bay National Park, mortality, primate conservation, roadkill, seasonality, wildlife-vehicle collisions, Zanzibar

## Abstract

Roads can affect wildlife in a variety of negative ways. Studies of road ecology have mostly concentrated in the northern hemisphere despite the potentially greater impact on biodiversity that roads may have in tropical habitats. Here, we examine a 4-year opportunistic dataset (January 2016 – December 2019) on mammalian roadkill observed along a road intersecting Jozani-Chwaka Bay National Park, Unguja, Zanzibar. In particular, we assess the impact of collisions on the population of an endangered and endemic primate, the Zanzibar red colobus (*Piliocolobus kirkii*). Primates accounted for the majority of roadkill. Monthly rainfall variation was not associated with roadkill frequency for mammals and specifically for the Zanzibar red colobus. No single age-sex class of colobus was found dead more often than expected given their availability in the local population. The exact effect of roadkill on colobus populations in habitats fragmented by roads is unknown given the lack of accurate, long-term life history data for this species. However, the frequency of kills documented in this study suggests further mitigation measures may be important. Our data show that mortality from collisions with vehicles in some groups of colobus are comparable to rates of mortality experienced by other primate populations from natural predation. Unlike natural predators, however, vehicles are not ‘selective’ in their targeting of ‘prey’. The long-term implications of such a ‘predation regime’ on this species remain to be established.

## Introduction

One of the many ways, in which roads affect wildlife is via increased mortality from vehicle-wildlife collisions (Tromblak & Frissell, 2000; Coffin, 2007; Newmark *et al.*, 2008; Laurance *et al.*, 2009). There are two key components to understanding the potential impact of vehicle collisions on the status of a species. First, we need to understand the scale of the problem by assessing the frequency of collisions and the factors determining the frequency, spatial and temporal distribution of roadkill. Second, we need to understand roadkill mortality effects on population structure and persistence.

Road ecology has made substantial progress in addressing the first issue. Numerous studies document patterns of wildlife-vehicle collisions across a variety of biomes (Rytwinski & Fahrig, 2012; Monge-Nájera, 2018; van der Ree *et al.*, 2015). Factors such as variation in rainfall can affect roadkill frequency with some studies showing more roadkill in wet, while others – in dry seasons (Jeganathan *et al.*, 2018; Njovu *et al.*, 2019). Roadkill are more frequent on roads near and within protected areas by comparison to roads outside them (Kioko *et al.*, 2015; Garrigia *et al.*, 2012; Akrim *et al*., 2019; Njovu *et al.*, 2019). Road type also has an effect as bigger, better roads, which allow for greater speed and larger vehicle access, cause greater mortality (Drews, 1995; Caro *et al.*, 2014; Epps *et al*. 2015; Collinson *et al*. 2019a). Finally, different species are not equally susceptible to road effects. Larger, longer-lived animals with slow life histories seem to be particularly vulnerable (Rytwinski & Fahrig, 2012).

Despite the abundant literature on roadkill, few studies have tackled the second issue, i.e., assessing the impact of road-induced mortality on population structure and persistence (e.g., Hels & Buchwald, 2001). Different individuals are at different risks of mortality. In Florida scrub jays, for example, the entire population along the roadside is at risk of roadkill mortality but those birds who immigrate into these areas are at higher risk within their first two years compared to other individuals. Similarly, fledglings near roads experienced significantly higher mortality in their first months of life but not later. This shows that individuals who find anthropogenic landscapes novel and struggle to navigate them safely are at higher risk (Mumme *et al.*, 2000). Differences in age and experience could explain roadkill mortality patterns in some mammals, also. Among groups of yellow baboons in Mikumi National Park, Tanzania, those that frequented a road intersecting the habitat, experienced average roadkill mortality of 10.3% of their population (Drews, 1995). Young, presumably less experienced baboons were, however, particularly susceptible: juvenile baboons were killed more frequently than expected based on their representation in groups that visited the road (*ibid*.). The need for further empirical data on this second question is urgent. Known effects of demographic mortality for populations show that the loss of a young reproductive female is more likely to have adverse effects on population maintenance than the loss of males or sexually immature individuals (Coulson *et al*., 2001). Demographic data on mortality patterns from roadkill are thus vital for predictive modelling of the impact of roads on endangered species, particularly slow reproducing ones, such as primates (Charnov & Berrigan, 2005). This issue for primates is further exacerbated when considering 60% of primate species are threatened by extinction and 75% have declining populations (Estrada *et al*., 2017). Given the limited literature on primate road ecology (Hetman *et al*., 2019) addressing this gap is of urgency.

Red colobus are one of the most endangered taxa of African primates (Struhsaker, 2005). The Zanzibar red colobus (*Piliocolobus kirkii*) is endemic to Unguja Island, Zanzibar. This species faces increasing pressure from habitat loss due to agriculture, human development and timber logging (Davenport, 2019). Although there are just over 5,800 individuals, according to the most recent and comprehensive census, the population seems to be in decline given its low recruitment rate by comparison to other red colobus taxa and primates more broadly (Davenport *et al*., 2019). Almost 50% of the Zanzibar red colobus population survive in the only national park in Zanzibar: Jozani-Chwaka Bay National Park (*ibid*.). They are the main attraction for visitors to the park and provide a crucial source of revenue for the local economy (Saunders, 2011; Carius & Job, 2019). While tourism brings important revenue, increased tourist traffic (in addition to other types of traffic) on the main road that intersects the southern edge of the national park poses a risk as colobus groups range on both sides of the road, cross it frequently and occasionally get hit by vehicles. In the absence of natural predators, roadkill incidents may be the most significant source of extrinsic mortality for this subpopulation of *P. kirkii*. Currently, however, there are no quantitative data to assess the magnitude of this problem for the colobus.

We describe a 4-year dataset on mammalian roadkill at Jozani-Chwaka Bay National Park and examine frequency and seasonality of reported deaths (carcasses that remained on the road). For mortality of Zanzibar red colobus in particular, we also examine the demographics of victims. We hypothesise that age-sex classes will be killed at different rates from their availability in the population of this primate. For sex differences we predict (1) adult males will be over-represented in the roadkill data, by comparison to adult females because of their potentially greater propensity for risk-taking. Alternatively, we predict that (2) adult females would be over-represented because they may be slower than males to cross, especially when carrying dependant offspring. Regarding age-class differences in road mortality, we predict that (3) subadults and juveniles would be over-represented in the roadkill because (a) they may be slower and lacking in learned abilities to avoid oncoming vehicles effectively, and perhaps (b) because subadults in particular would be the age-class most likely to disperse from their natal group. Alternatively, we predict that (4) adults would be over-represented in the roadkill dataset by comparison to immatures because adults may perceive roads as more familiar environments and so be less cautious when crossing.

## Methods

### Study site

Jozani-Chwaka Bay National Park (S6° 16.339’; E39° 25.154’) is located on Unguja Island, Zanzibar (Fig. 1A), and is the last stronghold of the endemic Zanzibar red colobus (Siex, 2003; Davenport *et al*., 2019). At its southern edge, a major road (approximately 5 m wide) intersects the home ranges of multiple colobus groups (Fig. 1B). These groups often cross the road in search of food and sleeping sites, creating the risk of road collisions and fatalities. There are four speed bumps along the road spanning approx. 600 m but colobus groups often range and cross the road beyond this speed buffer zone.

**Figure 1.**
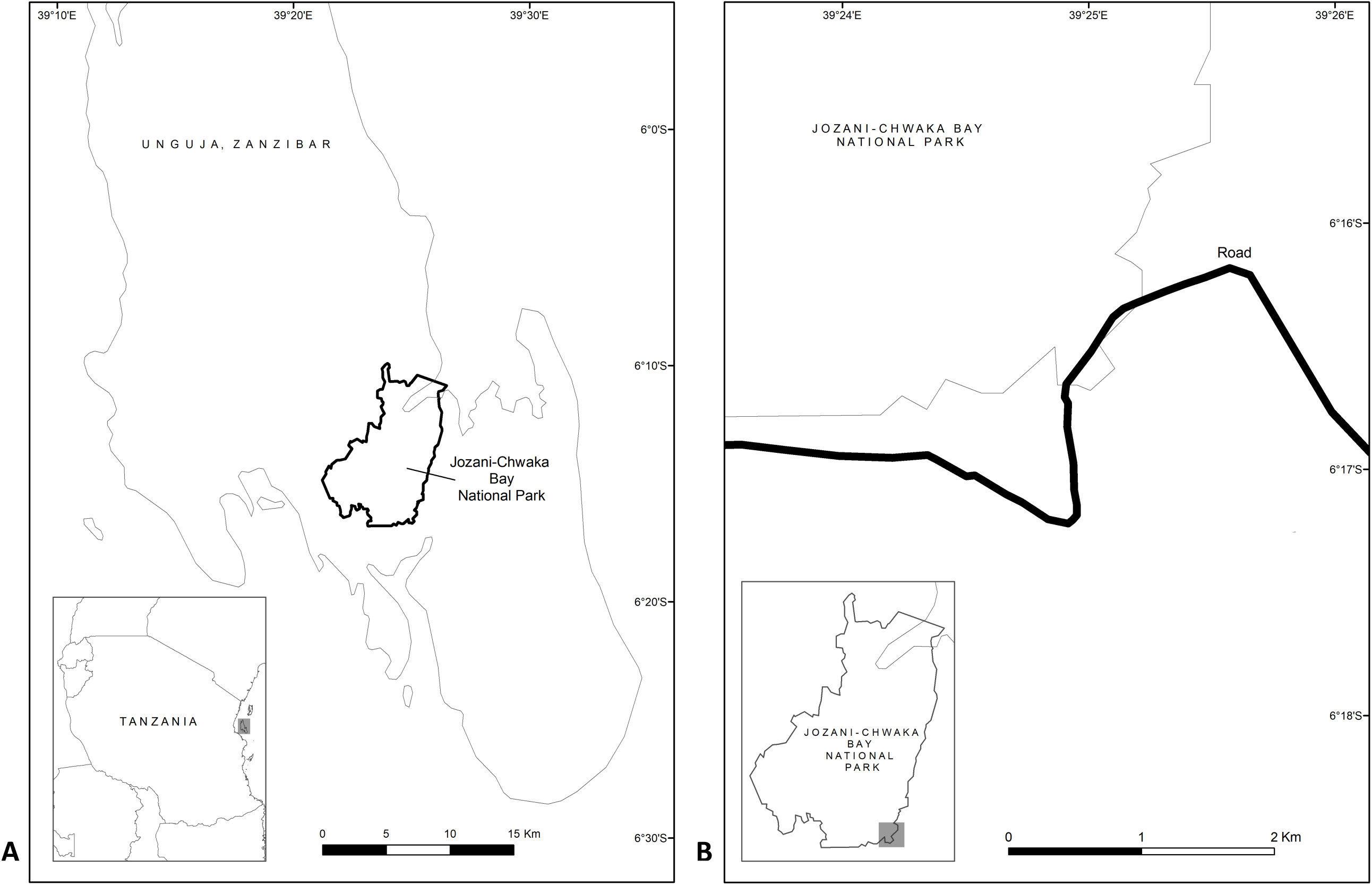
Location of study site: (A) Jozani-Chwaka Bay National Park, Zanzibar, Tanzania; (B) A view of the road from which roadkill data were collected.

### Roadkill data

Roadkill was recorded opportunistically by Jozani-Chwaka Bay National Park staff along the main road intersecting the park at its southern edge over 4 years based both on direct observation and on reports from the public (January 2016 – December 2019; 43 months of records were available; 5 months missing). Though no systematic road surveys were conducted, all staff members working at park headquarters commuted from nearby villages into work on a daily basis along the main road. This allowed them to spot roadkill (though probably paying more attention to colobus). Staff also monitored the road nearest to headquarters repeatedly during the day as they led groups of tourists around the area in search of colobus on both sides of the road. Dead colobus further away from headquarters are reported voluntarily and without reward by members of the public to park staff. Based on descriptions of approximate locations in the notes, we estimate that roadkill was reported from a section of the road between 3.4 km and 6.54 km in length. While all mammals were counted, if found or reported, we assume that indirect reports (by members of the public) have mostly focused on Zanzibar red colobus (rather than on smaller mammals). In most cases (for colobus), in addition to the date of observation, the age class and sex of dead monkeys was also noted. Records of individuals who may have been hit but did not die on the road were not available.

### Rainfall data

To examine the association between roadkill frequency and monthly rainfall, we obtained daily precipitation data for Jozani-Chwaka Bay National Park for January 2016 - December 2019 (43 months) from the NASA Langley Research Center (LaRC) POWER Project funded through the NASA Earth Science/Applied Science Program.

### Demographic composition of roadside colobus groups

To examine if colobus of different age-sex classes were equally like to die on the road as a result of vehicle collision or if some age-sex classes were at greater risk, we compared the representation of these classes in the roadkill dataset to that in a census of groups near the road. Data on group composition come from a detailed census conducted in the Jozani-Chwaka Bay NP area in 2014 and described in detail by Davenport *et al.* (2019). Briefly, survey teams located colobus via a total sweep census and followed each group for 2 – 3 days to count all of its members and obtain data on its movement. Age-sex classes were identified by trained and experienced observers with a high degree of inter-observer reliability (Davenport *et al.* 2019).

Based on the spatial data recorded during these group follows in 2014 and our knowledge of the distribution and movements of groups in the same area in 2018 – 2019 we selected 18 groups from the total of 151 groups identified in the JCBNP area by survey teams (Davenport *et al.*, 2019) that met our definitions of being ‘roadside groups’. We considered ‘roadside groups’ to be those which were observed within a 300 m perpendicular distance from the road on at least one occasion during the group follow. From these 18 groups, six were seen to cross the road and one came within 5 m of the road bit did not cross it. For the remaining 11 groups, the mean minimum perpendicular distance to the road at which they were observed was 94.8 m (range: 17 – 217 m). The mean furthest perpendicular distance from the road that these same 11 groups were observed was 378.2 m (range: 221 – 590 m). Although the amount of ranging data for each of the groups was limited (52 group follow days; two groups followed for two consecutive days; 16 for three), given our knowledge of the area and of Zanzibar red colobus ranging ecology, we included all 18 groups in analysis as ‘roadside groups’, a reasonable approximation of their potential likelihood of crossing the road. Groups that were found further than 300 m from the road, while technically capable of crossing are not likely to do so often, given the presence of other groups’ home ranges between them and the road. Although Zanzibar red colobus are not strictly territorial and they exhibit substantial intergroup home range overlap (Siex, 2003; Struhsaker, 2010), based on observations in 8 groups ranging at various distances from the road over a 15 month period (Zanzibar Red Colobus Project, unpublished data) we feel reasonably confident our 300 m cut-off did not exclude any groups that were frequent road-crossers. If anything, this cut-off is likely a liberal assumption of road crossing likelihood. Thus, in addition to using the 18 groups as the baseline population to estimate vehicle collision mortality rates, we considered two more conservative definitions of ‘roadside groups’ in our analyses. First, we restricted the comparison to only the 13 groups that were seen within 100 m of the road, and then restricted the demographic dataset further to only those seven groups that ranged closest to the it (i.e., the six that were seen to cross it and the one that was seen within 5 m of it). Restricting the original 18-group dataset to the 13 or the 7 groups nearest the road did not affect the statistical significance of the results.

### Statistical analyses

We analysed data in R v. 3.6.1 (R Core Team, 2018). We tested the association between rainfall and roadkill frequencies with the Pearson correlation test. We compared the age-sex class composition of roadkill to the demography of the local population of colobus via Fisher’s exact test (for sex class) and the likelihood ratio test across the different age classes (package “MASS”: Venables & Ripley, 2002).

## Results

Over the 4-year study period (43 months of opportunistic data collection) we recorded 7 different mammal species as roadkill (Table 1). The majority (66%) of carcasses were of diurnally active animals. Primates were most at risk: they accounted for 83% of all recorded deaths. The endemic and threatened Zanzibar red colobus were particularly common in the dataset with 55% of all carcasses.

**Table 1.** Roadkill recorded along the main road near Jozani-Chwaka Bay National Park, Zanzibar (January 2016 – December 2019; N = 43 months of recording).

| Species | Activity pattern | Total roadkill | Kills/month |
| --- | --- | --- | --- |
| Zanzibar red colobus (*Piliocolobus kirkii*) | Diurnal | 29 | 0.67 |
| Small-eared greater galago (*Otolemur garnetti*) | Nocturnal | 13 | 0.30 |
| Rats (Muridae) | Nocturnal | 3 | 0.07 |
| Bushy-tailed mongoose (*Bdeogale crassicauda*) | Nocturnal | 2 | 0.05 |
| Black and rufous elephant shrew (*Rhynchocyan petersi*) | Diurnal | 2 | 0.05 |
| White-collared guenon (*Cercopithecus albogularis*) | Diurnal | 2 | 0.05 |
| Squirrels (Sciuridae) | Diurnal | 2 | 0.05 |
| **Total** |  | **53** | **1.23** |

On average, there were 1.23 carcasses found on the road per month (range: 0 – 6). Rainfall at the study site had a mean of 105 mm (range: 5 – 456 mm). Variance in rainfall, however, was unrelated to all-species roadkill frequency (*r*_s_ = −0.248, N = 43 months, P = 0.109, Fig. 2).

**Figure 2.**
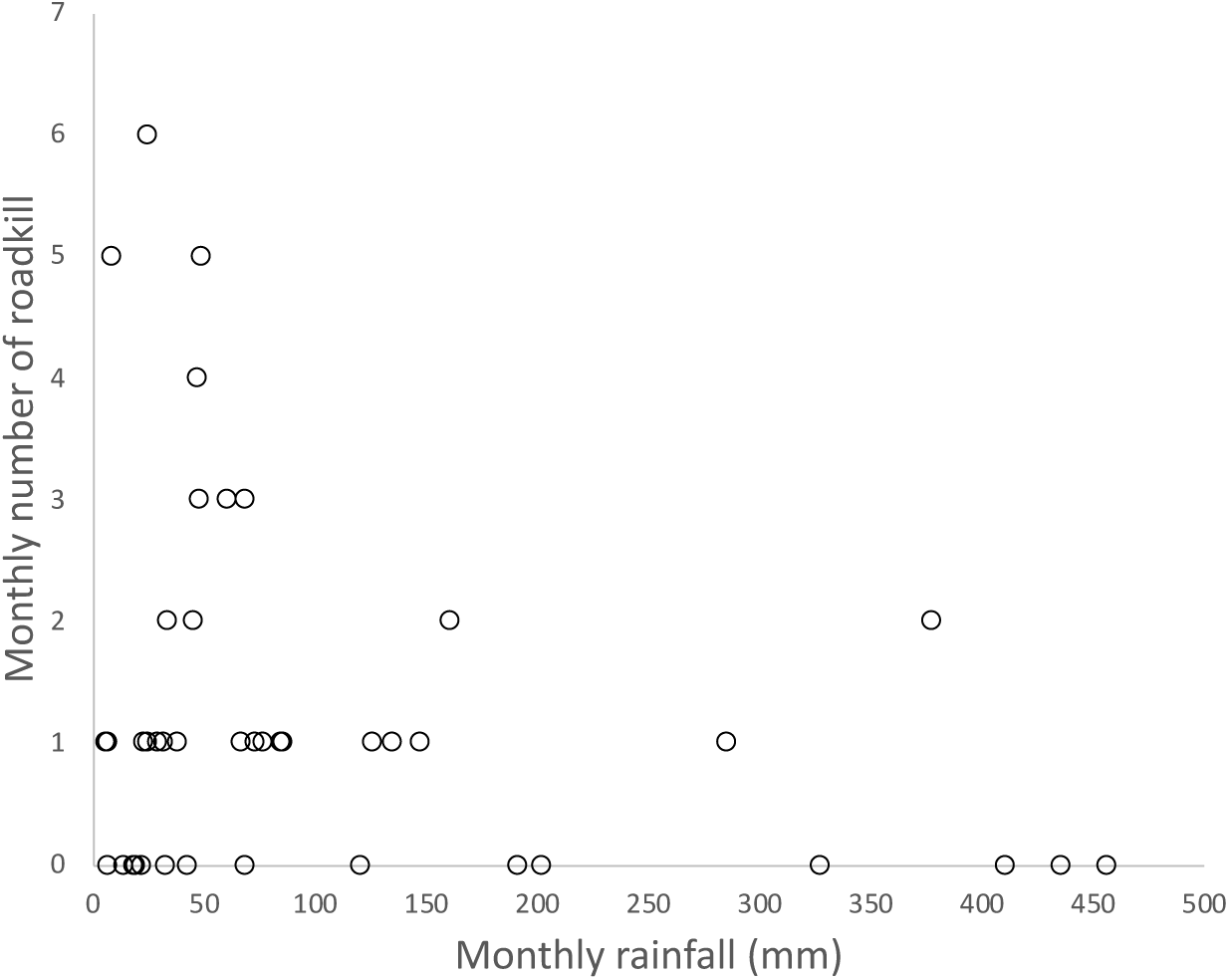
Relationship between monthly rainfall at Jozani-Chwaka Bay National Park and mammalian roadkill frequency along the Jozani road (N = 43 months).

For Zanzibar red colobus, specifically, there were a mean of 0.67 carcasses found on the road per month (range: 0 – 6 deaths). Rainfall was not related to the monthly frequency of colobus roadkill either (*r*_s_ = −0.128, N = 43 months, P = 0.412). The ratio of adult female to male colobus in the roadkill dataset was 2.75, while in the 18 roadside groups it was 3.54 (Table 2). However, the relative proportion of males and females that were killed did not differ significantly from their numbers in the roadside groups (Fisher’s exact test: odds ratio = 1.287, P = 0.750; Fig. 3A). The proportions of different age classes in the roadkill dataset did not differ from their expected values based on the demographic composition of the 18 roadside groups either (Likelihood Ratio test: χ^2^= 5.302, df = 2, P = 0.071; Fig. 3B & Table 2). Different age-sex classes were therefore equally likely to be found dead on the road, given their availability in the local population. This result did not change across the different definitions of ‘roadside groups’ we considered (Table 2).

**Table 2.** Annual mortality rate (%) of Zanzibar red colobus caused by vehicle collisions on the road intersecting Jozani-Chwaka Bay National Park. Different mortality rate estimates are presented based on three empirically-derived levels of confidence as to which colobus groups are directly exposed to the risk of collision (see *Methods* for defining the three populations). Note that the sum of adult females and adult males does not equal the number of adults in the roadkill data, as some of the adults killed were not sexed in the records.

|  |  | Population estimates for groups at various distances from the road |  |  | Annual mortality % to vehicle collisions |  |  |
| --- | --- | --- | --- | --- | --- | --- | --- |
| Age-sex class | Roadkill observed | 18 groups (300 m roadside) | 13 groups (100 m roadside) | 7 groups (observed crossing) | 18 groups (300 m roadside) | 13 groups (100 m roadside) | 7 groups (observed crossing) |
| *Adults by sex* |  |  |  |  |  |  |  |
| Adult females | 11 | 209 | 167 | 106 | 1.32 | 1.65 | 2.59 |
| Adult males | 4 | 59 | 48 | 32 | 1.69 | 2.08 | 3.13 |
| *All individuals by age* |  |  |  |  |  |  |  |
| All adults | 19 | 268 | 215 | 138 | 1.77 | 2.21 | 3.44 |
| Subadult/juvenile | 8 | 61 | 50 | 39 | 3.28 | 4 | 5.13 |
| Infants | 2 | 80 | 69 | 47 | 0.63 | 0.72 | 1.06 |
| *Total* | 29 | 409 | 334 | 224 | 1.77 | 2.17 | 3.24 |

**Figure 3.**
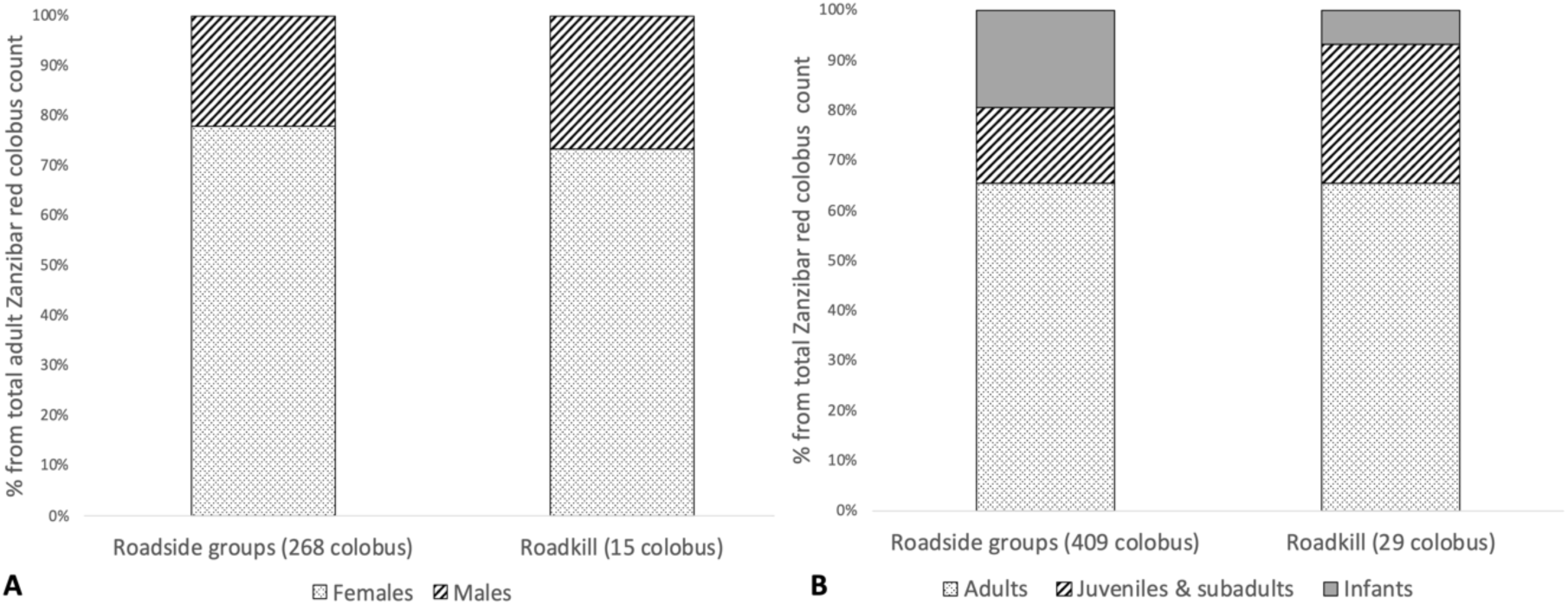
Composition of the roadkill dataset for Zanzibar red colobus by (A) sex (N = 15 adults of known sex); and by (B) age class (29 colobus), in comparison to the demographics of the 18 roadside groups (268 adults and 409 total colobus, respectively). No significant differences were found (see text for details).

To consider the overall impact of vehicle collisions on population-level mortality, we estimated annual mortality rates using the three ‘subpopulations’ that we considered at risk (depending on the confidence in our definition of their likelihood of crossing the road). If only the 7 groups nearest the road are considered, their annual loss to collisions is at 3.24%. This number is lower if we consider all 18 groups within about 300 m of the road to be at equal risk of collisions (Table 2).

## Discussion

This is the first study to quantify the effects of vehicle collisions on mortality in the Zanzibar red colobus, an endangered endemic primate. This dataset is an important contribution to the road ecology literature more broadly, given the lower number of roadkill studies in the tropics, by comparison to the northern hemisphere (Monge-Nájera, 2018; van der Ree *et al*., 2015; Collinson *et al.* 2019a). Primates, and colobus, in particular, were the most common species found as roadkill. It is likely that the absolute number of non-colobus deaths, however, is an underestimate. The national park and the tourism industry in the area are focused on this charismatic species of primate, so other animals that die on the road may go unreported by members of the public. Our finding that diurnal species were more often found on the road, contradicts previous findings from Africa which found a higher number of nocturnal roadkill (Kioko, *et al*., 2015; Njovu, *et al*., 2019). This could be a reflection of the site differences in habitats and species diversity between studies but could also reflect methodological differences. Many previous studies relied exclusively on early-morning road surveys to quantify roadkill, whereas our opportunistic records were not constrained to the early hours of the day. Up to 62% of carcasses on roads can be removed by scavengers in a period as short as two hours (Schwartz *et al.*, 2018) so animals that die during the day may not remain there until the next road survey the following morning.

Monthly variation in rainfall was not related to monthly roadkill frequency at Jozani, for neither all mammals or for colobus separately. This suggests that, unlike in some other study systems (e.g., da Rosa & Bager, 2012; Machado *et al*., 2015; Jeganathan *et al*, 2018; Njovu *et al*., 2019), precipitation may not strongly influence mammal movements in the Jozani area or the behaviour of drivers on the road. Alternatively, it may be that our dataset underestimates roadkill frequency during the wet months of the year. As the records of roadkill were collected opportunistically by national park staff and/or reported by members of the public, it is feasible such monitoring was less extensive under inclement weather. Future systematic roadkill surveys would resolve this.

Contrary to our predictions, males and females were found dead on the road in similar proportions to their availability within surrounding roadside groups. Likewise, adults, subadults/juveniles and infants were observed as roadkill as often as expected. This suggests that the way colobus cross the roads does not place any particular age-sex class at a significant disadvantage. Behavioural data on individual rates of crossing and collision risk would allow further tests of this hypothesis. The estimates of group demographics we used to calculate the mortality rates for colobus surrounding the road may be biased in two important ways. First, data on group composition were collected in 2014, while data on roadkill come from 2016 – 2019. Second, in the demographic data from 2014 we did not distinguish subadults from juveniles. A similarly ambiguous classification of immature animals applies to the roadkill records. This is therefore an imperfect but still valuable preliminary analysis, that would serve as a baseline for future more detailed surveys of roadkill mortality in this species.

We have estimated the annual loss of colobus to vehicle collisions to be in the range of 1.77 – 3.24% (depending on the baseline estimate for the population which is at risk; Table 2). To put this figure in context, data on mortality rates from natural predators could be informative. The Zanzibar red colobus experience very limited natural predation at Jozani. The Zanzibar leopard is likely extinct, there are no crowned eagles, and pythons are uncommon. The only predation this species experiences is from feral and village dogs (Georgiev *et al.*, 2019) and people (Davenport *et al.*, 2019), though reliable data for estimating the magnitude of these sources of mortality are not available. Data on natural predation on primates from the wild, in general, are difficult to obtain and are available from only a few long-term studies of individually recognised primates. One review of primate predation places an estimate of annual mortality from predation in the range of 0 – 15% (Cheney & Wrangham, 1987). While more recent studies have shown that under exceptional circumstances predation rates can be even higher (e.g., 40 −50% of red colobus killed by chimpanzees in Ngogo, Uganda: Teelen, 2008), mean predation rate across species, populations and habitats can be estimated at around 6% (Cheney & Wrangham, 1987). Our most conservative estimate of 3.24% annual road mortality for the Zanzibar red colobus, therefore, falls within the lower range of mortality rates experienced by different primates from natural predation. This rate of mortality is, in fact, higher than natural predation mortality for some red colobus populations elsewhere. For example, at Mahale, Tanzania in some years annual predation loss has been estimated at only 1.1 – 1.3% (Boesch *et al.*, 2002). Unlike natural predators, which are more successful in killing animals in poorer condition, vehicles remove individuals from a population without regard to their condition or health (Bujoczek *et al.*, 2011). Even non-lethal collisions can have a negative impact on animals via increases in glucocorticoid production (Narayan, 2019). Taken together, the direct and indirect effects of colobus-vehicle collisions could lead to suboptimal population health that would contribute to reduced persistence of roadside groups.

In line with observations elsewhere (e.g., Drews, 1995) after the Jozani road was tarmacked in 1996, vehicles started travelling at greater speed and roadkill became a more common problem (Struhsaker & Siex, 1998). National park staff at the time suggested that perhaps a colobus was killed every 2 – 3 weeks (*ibid*.). Struhsaker & Siex (1998) estimated, based on a population of 150 colobus exposed to the road, that 12 – 17% of colobus were lost to road accidents every year. By comparison, our data show lower mortality in 2016 – 2019: almost 1 colobus died every 6 weeks and we estimate an annual mortality loss in the range of 1.77 – 3.24% (Table 2). Changes in population size and group ranging patterns in the 20 years that separate these estimates may account for the difference in mortality rates. A major factor, however, has been the installation of four speed bumps and colobus-crossing warning signs near the main entrance of the national park, which reportedly achieved an 80% reduction in mortality in the years after they were built (Thomas Struhsaker, pers. comm. in Epps *et al.*, 2015). As these speed bumps only cover about 600 m of the road that intersects colobus ranges, extending the speed bump zone for the entirety of the road that lies within the known range of colobus near the road (approx. 1.8 km), would likely bring further reductions in roadkill.

Measures that are unlikely to be ignored by drivers (e.g., speed bumps) have a bigger effect on reducing roadkill and are the most effective solution (Rytwinski *et al.* 2016). But complementing this approach with additional, less costly, interventions can also be beneficial. Warning signs containing photographic images of animals placed near roadkill hotspots can bring about additional reductions in roadkill by altering driver behaviour over short distances (Collinson *et al.* 2019b). Such mitigation measures however can become less effective as drivers habituate to them but can still be effective for visitors (Huijser *et al*., 2015). Updating the roadside signage more frequently with visually captivating images of colobus in their natural habitat may serve the dual purpose of both preventing driver habituation to these warning signs and attracting more tourists to the area by highlighting the ecotourism opportunity. Some colobus groups elsewhere on the island also inhabit areas where roads intersect their home ranges (Fig 1 in Davenport *et al.* 2019). Identifying roadkill hotspots outside the immediate area of Jozani-Chwaka Bay National Park and carrying out road mitigation may provide additional benefits for slowing the decline of this primate throughout its range.

Road networks in Africa are projected to increase by 60% between 2000-2050 (Dulac, 2013). The continued growth of the tourism industry with its demand for quick, easy and efficient travel (Khadaroo & Seetanah, 2008) will worsen the already serious issues of wildlife-vehicle collisions (Caro *et al*., 2014). Understanding the impact of roads on animal populations and the most effective way to counteract them will be crucial in achieving a balance between the demands of economic development and wildlife conservation.

## Author contributions

Study design and fieldwork: HO, MKM, AJM, TRBD, AVG; data analysis: HO, MELG, AVG; wrote the first draft: HO & AVG; comments and revisions: HO, MKM, AJM, MELG, TRBD, AVG.

## Acknowledgements

We thank the Department of Forestry and Non-Renewable Natural Resources of the Revolutionary Government of Zanzibar for permission to conduct this research. Ali Kassim, Ameir Abdalla, and Mwinyi Abdalla provided field assistance in data collection. We also thank Kassim Madeweya and Said Fakih for logistical advice and assistance. Zoe Melvin, John Healey and Carlos Cardenas-Iniguez provided helpful comments on this work. The Zanzibar Red Colobus Project was supported by funding to AG from the Royal Society and Bangor University.

## Conflicts of interest

None.

## Ethical standards

This research was approved by the Ethics Committee of the College of Environmental Sciences and Engineering of Bangor University. Research permission was obtained from the Department of Forestry and Non-Renewable Natural Resources of the Revolutionary Government of Zanzibar.

## References

Akrim, F., Mahmood, T., Andleeb, S., Hussain, R. & Collinson, W.J. (2019) Spatiotemporal patterns of wildlife road mortality in the Pothwar Plateau, Pakistan. Mammalia, 83(5).

Boesch, C., Hohmann, G. & Marchant, L. (eds) (2002) Behavioural Diversity in Chimpanzees and Bonobos. Cambridge University Press, Cambridge, UK.

Bujoczek, M., Ciach, M. & Yosef, R. (2011) Road-kills affect avian population quality. Biological Conservation, 144(3), 1036–1039.

Carius, F. & Job, H. (2019) Community involvement and tourism revenue sharing as contributing factors to the UN Sustainable Development Goals in Jozani–Chwaka Bay National Park and Biosphere Reserve, Zanzibar, Journal of Sustainable Tourism, 27(6), 826–846

Caro, T., Dobson, A., Marshall, A.J. & Peres, C.A. (2014) Compromise solutions between conservation and road building in the tropics. Current Biology, 24(16), R722–R725.

Charnov, E. & Berrigan, D. (2005) Why do female primates have such long lifespans and so few babies? or Life in the slow lane. Evolutionary Anthropology: Issues, News, and Reviews, 1(6), 191–194.

Cheney, D.L. & Wrangham, R.W. (1987) Predation. In Primate Societies (eds. Smuts, B.B., Cheney, D.L., Seyfarth, R.M., Wrangham, R.W. & Struhsaker, T.T.). University of Chicago Press, Chicago, USA.

Coffin, A.W. (2007) From roadkill to road ecology: A review of the ecological effects of roads. Journal of Transport Geography, 15(5), 396–406.

Collinson, W.J., Davies-Mosert, H., Roxburgh, L. & van der Ree, R. (2019a) Status of road ecology research in Africa: Do we understand the impacts of roads, and how to successfully mitigate them? Frontiers in Ecology and Evolution, 7(479).

Collinson, W.J., Marneweck, C. and Davies-Mostert, H.T. (2019b). Protecting the protected: reducing wildlife roadkill in protected areas. Animal Conservation. 22(4): 396–403.

Coulson, T., Catchpole, E.A., Albon, S., Morgan, B., Pemberton, J.M., Clutton-Brock, T.H. et al. (2001) Age, sex, density, winter weather, and population crashes in Soay sheep. Science, 292(5521), 1528–1531.

da Rosa, C.A. & Bager, A. (2012) Seasonality and habitat types affect roadkill of neotropical birds. Journal of Environmental Management, 97(1), 1–5.

Davenport, T. (2019) Piliocolobus kirkii. In The IUCN Red List of Threatened Species 2019: e.T39992A92630664. https://dx.doi.org/10.2305/IUCN.UK.2019-3.RLTS.T39992A92630664.en. [Downloaded on 09 January 2020].

Davenport, T., Fakih, S., Kimiti, S., Kleine, L., Foley, L., & De Luca, D. (2019) Zanzibar’s endemic red colobus *Piliocolobus kirkii*: First systematic and total assessment of population, demography and distribution. Oryx, 53(1), 36–44. doi:10.1017/S003060531700148X

Drews, C. (1995) Road kills of animals by public traffic in Mikumi National Park, Tanzania, with notes on baboon mortality. African Journal of Ecology, 33(1).

Dulac, J. (2013) Global land transport infrastructure requirements. Paris: International Energy Agency, p.20.

Epps, C.W., Nowak, K. & Mutayoba, B. (2015) Unfenced reserves, unparalleled biodiversity and a rapidly changing landscape: roadways and wildlife in East Africa. In Handbook of road ecology (eds. Van Der Ree, R., Smith, D.J. & Grilo, C.), pp.448–454. John Wiley & Sons Inc., West Sussex, UK.

Estrada, A., Garber, P.A., Rylands, A.B., Roos, C., Fernandez-Duque, E., Di Diore, A. et al. (2017) Impending extinction crisis of the world’s primates: Why primates matter. Sciences Advances, 3(e:1600946).

Garrigia, N., Santos, X., Montori, A., Richter-Boix, A., Franch, M. & Llorente, G. (2012) Are protected areas truly protected? The impact of road traffic on vertebrate fauna. Biodiversity and Conservation, 21, 2761–2774.

Georgiev, A., Melvin, Z., Warkentin, A., Winder, I. & Kassim, A. (2019) Two cases of dead-infant carrying by female Zanzibar red colobus (*Piliocolobus kirkii*) at Jozani-Chwaka Bay National Park, Zanzibar. African Primates, 13, 57–60.

Hels, T. & Buchwald, E. (2001) The effect of road kills on amphibian populations. Biological conservation, 99(3), 331–340.

Hetman, M., Kubicka, A., Sparks, T. & Tryjanowski, P. (2019) Road kills of non-human primates: a global view using a different type of data. Mammal Review, 49(3), 276–283.

Huijser, M.P., Mosler-Berger, C., Olsson, M. & Strein, M. (2015) Wildlife warning signs and animal detection systems aimed at reducing wildlife-vehicle collisions. In Handbook of road ecology (eds. Van Der Ree, R., Smith, D.J. & Grilo, C.) pp.198–212. John Wiley & Sons Inc., West Sussex, UK.

Jeganathan, P., Mudappa, D., Kumar, M.A. & Shankar Raman, T.R. (2018) Seasonal variation in wildlife roadkills in plantations and tropical rainforest in the Anamalai Hills, Western Ghats, India. Current Biology, 114(3), 619–626.

Khadaroo, J. & Seetanah, B. (2008) The role of transport infrastructure in international tourism development: A gravity model approach. Tourism Management, 29(5), 831–840.

Kioko, J., Kiffner, C., Jenkins, N. & Collinson, W. (2015) Wildlife roadkill patterns on a major highway in northern Tanzania. African Zoology, 50(1), 17–22.

Laurance, W.E., Goosem, M. and Laurance, S.G. (2009) Impacts of roads and linear clearings on tropical forests. Trends in Ecology & Evolution, 24(12), 659–669.

Machado, F., Fontes M., Mendes, P., Moura, A. & Romao, B. (2015) Roadkill on vertebrates in Brazil: seasonal variation and road type comparison. North-Western Journal of Zoology, 11(2), 247–252.

Monge-Nájera, J. (2018) Road kills in tropical ecosystems: A review with recommendations for mitigation and for new research. Revista de biologia tropical, 66(2), 722–738.

Mumme, R.L., Schoech, S.J., Woolfenden, G.E. & Fitzpatrick, J.W. (2000) Life and death in the fast lane: Demographic consequences of road mortality in the Florida scrub-jay. Conservation Biology, 14(1), 501–512.

Narayan, E. (2019) Physiological stress levels in wild koala sub-populations facing anthropogenic induced environmental trauma and disease. Scientific Reports, 9, 6031. doi:10.1038/s41598-019-42448-8

Newmark, W.D. (2008) Isolation of African protected areas. Frontiers in Ecology and the Environment, 6(6).

Njovu, H., Kisingo, A., Hesselberg, T. & Eustace, A. (2019) The spatial and temporal distribution of mammal roadkills in the Kwakuchinja Wildlife Corridor in Tanzania. African Journal of Ecology, 57(3), 423–428.

R Core Team (2018) R: A language and environment for statistical computing. R Foundation for Statistical Computing, Vienna, Austria. URL: https://www.R-project.org/.

Rytwinski, T. & Fahrig, L. (2012) Do species life history traits explain population responses to roads? A meta-analysis. Biological Conservation, 147(1), 87–98.

Rytwinski, T., Soanes, K., Jaeger, J.A., Fahrig, L., Findlay, C.S., Houlahan, J., et al. (2016) How effective is road mitigation at reducing road-kill? A meta-analysis. PLoS one, 11(11), p. e0166941.

Saunders, F. (2011) It’s like herding monkeys into a conservation enclosure: The formation and establishment of the Jozani-Chwaka Bay National Park, Zanzibar, Tanzania. Conservation and Society, 9(4), 261–273.

Schwartz, A.L.W., Williams, H.F., Chadwick, E., Thomas, R.J. & Perkins, S.E. (2018) Roadkill scavenging behaviour in an urban environment. Journal of Urban Ecology, 4(1).

Siex, K.S. (2003) Effects of population compression on the demongraphy, ecology, and behaviour of the Zanzibar red colobus monkey (Procolobus kirkii). PhD thesis, Duke University, Durham, North Carolina, USA.

Struhsaker, T.T. (2005) Conservation of red colobus and their habitats. International Journal of Primatology, 26(3), 525–538.

Struhsaker, T.T. (2010) The Red Colobus Monkeys: Variation in Demongraphy, Behavious, and Ecology of Endangered Species. Oxford University Press, Oxford, UK.

Struhsaker, T.T. & Siex, K.S. (1998) The Zanzibar red colobus monkey: Conservation status of an endangered island endemic. Primate Conservation, 18, 51–58.

Teelen, S. (2008) Influence of chimpanzee predation on the red colobus population at Ngogo, Kibale National Park, Uganda. Primates, 49(1), 41–9.

Trombulak, S.C. & Frissell, C.A. (2000) Review of ecological effects of roads on terrestrial and aquatic communities. Conservation Biology, 14, 18–30.

van der Ree, R., Smith, D.J. & Grilo, C. (eds.) (2015) Handbook of road ecology. John Wiley & Sons Inc., West Sussex, UK.

Venables, W.N. & Ripley, B.D. (2002) Modern applied statistics with S (4^th^ Edition). Springer, New York, USA.

